# Okapi-EM – a napari plugin for processing and analysing cryogenic serial FIB/SEM images

**DOI:** 10.1101/2022.12.15.520541

**Authors:** Luís M. A. Perdigão, Elaine M. L. Ho, Zhiyuan C. Cheng, Neville B.-y. Yee, Thomas Glen, Liang Wu, Michael Grange, Maud Dumoux, Mark Basham, Michele C. Darrow

## Abstract

An emergent volume electron microscopy (vEM) technique called cryogenic serial plasma focused ion beam milling scanning electron microscopy (pFIB/SEM) can decipher complex biological structures by building a three-dimensional picture of biological samples at mesoscale resolution. This is achieved by collecting consecutive SEM images after successive rounds of FIB milling that expose a new surface after each milling step. Due to instrumental limitations, some image processing is necessary before 3D visualisation and analysis of the data is possible. SEM images are affected by noise, drift, and charging effects, that can make precise 3D reconstruction of biological features difficult. This paper presents Okapi-EM, an open-source Napari plugin^(1)^ developed to process and analyse cryogenic serial FIB/SEM images. Okapi-EM enables automated image registration of slices, evaluation of image quality metrics specific to FIB-SEM imaging, and mitigation of charging artefacts. Implementation of Okapi-EM within the Napari framework ensures that the tools are both user- and developer-friendly, through provision of a graphical user interface and access to Python programming. Napari also hosts a variety of other image processing plugins so Okapi-EM tools can be integrated into and combined with other workflows. Okapi-EM can be downloaded freely at https://github.com/rosalindfranklininstitute/okapi-em, or installed from Python package index (PyPI).

**Impact statement:** Cryogenic serial pFIB/SEM is an emerging microscopy technique that is used to visualise 3D structures of biological features at mesoscale resolutions^(2)^. This technique requires common post processing of data such as alignment and charge mitigation to enable robust segmentation and analysis. In addition, approaches are needed to quantify data quality to enable an assessment of features and tune data acquisition parameters to enable optimal image acquisition. This article presents Okapi-EM, a combination of software tools designed to facilitate these important initial steps in assessing and processing images from these experiments. These tools have been assembled as a plugin for a popular 3D biological image visualiser called Napari, making their usage user-friendly and readily accessible.

## Introduction

Recent advances in cryo-electron microscopy hardware have seen an emergence of instruments which combine microscopy techniques and milling instruments within the same system. Dual-beam focused ion beam scanning electron microscopy (FIB/SEM), maintained at cryogenic temperatures, provides a workflow to acquire volumetric SEM images of a range of biological samples in their near native state at nanometre-resolution^(2–4)^. This technique builds volumetric representation of the specimen by cyclic FIB-milling (to remove the freshly imaged surface) and SEM imaging of the specimen. Several SEM images (typically hundreds) are obtained, corresponding to decreasing heights of the specimen, which can be computationally stacked to obtain a volumetric representation of the sample being measured.

Serial FIB/SEM has historically been used to capture images of fixed, stained and resin-embedded samples providing cellular and sub-cellular imaging in 3D that can be used for the reconstruction and analysis of biological features. However, fixation, staining and resin-embedding introduce artefacts^(5–7)^. With the development of cryogenically stable stages and reduced rates of ice formation it is possible to image cryogenically prepared samples providing structural information of samples in near native states. Cryogenic serial FIB/SEM has been done using modified room temperature plasma ion beam milling microscopes which were designed for physical science applications and more recently has been further enabled through the development of fit-for-purpose commercial options, some of which offer multiple plasma generation gases^(2,8–13)^.

As with many experimental techniques which produce 3D data, it is often desirable to annotate biological features and to visualise the structure in three-dimensions (3D), but important pre-processing steps are needed before data is suitable as input for these segmentation tasks. For instance, small translational movements between images within the stack caused by stage and/or sample movement are often observed as misalignment between images; while compensatory functions exist within the instrument it is not always possible to correct on-the-fly. Therefore, these must be compensated for, otherwise volumetric segmentation ^(14,15)^ and computational counting tools like connected components algorithms will fail or struggle to succeed. Additionally, SEM images of biological samples often contain artefacts caused by charging around insulating substances such as lipids^(4,11)^. Automatic and semi-automatic segmentation tools require aligned datasets and data that can be effectively normalised to remove any strong features generated by sample charging. Another common issue observed during FIB/SEM imaging is the creation of curtaining artefacts during the milling step which are then visible as streaks in the milling direction during SEM imaging. Finally, having quantitative tools that assess the quality of the data under certain imaging (e.g., optimal focus, voltage, etc) and milling conditions (e.g., focusing, curtaining, milling accuracy) will assist in the generation of data that best mitigate these factors, resulting in optimal further processing and optimisation of future data acquisition strategies^(14,15)^. These necessary pre-segmentation tasks can be time-consuming and often require use of multiple pieces of software or bespoke code. Okapi-EM provides a selection of tools which address some of these needs in a single software package found within Napari^(1)^. In this first release of Okapi-EM, there are three tools available:

1. Stack Alignment. This tool provides the user with appropriate transformation options for alignment of stacks of slices.
2. Charge mitigation (Chafer). This tool requires pre-segmented “charge centres”, then applies filters to mitigate the charging artefacts found nearby.
3. Resolution estimation (Quoll). This tool requires microscope calibration and provides a measure of the mean resolution and standard deviation for individual slices.

Example data used throughout this manuscript are available on EMPIAR and their sample preparation and FIB/SEM data collection has been described in Dumoux et al.^(2)^.

### Implementation

Napari is an open-source, user-friendly, data visualisation application that runs in Python programming language^(1)^. It supports three-dimensional visualisation of different data types (i.e., image, labels) interactively. Its annotation features are useful in biological image analysis and processing. Furthermore, its functionality can be extended with support for plugins, allowing continuous development of image processing methods alongside of experimental microscopy advances. Developing within the Python ecosystem also facilitates deep learning approaches with easy access to modern machine learning packages such as scikit-learn^(16)^ and PyTorch^(17)^. The plugin installation is uniquely easy as it uses the well-established package management system (PyPI), which also enables plugin installation chains, where a single plugin can install many other plugins or packages as needed to create a *bundle*. Bundling several plugins in this way means that a whole toolbox can be created for specific data processing workflows or research themes (e.g., devbio-napari^(18)^, or napari-assistant^(19)^, both of which were tailored for biological image processing). Currently Okapi-EM incorporates three major image processing tools which are organised in the user interface as separate tabs: stack-alignment, charge suppression and resolution measurement (Figure 1). Each tab contains user interface elements to adjust settings and execute data analysis.

**Figure 1.**
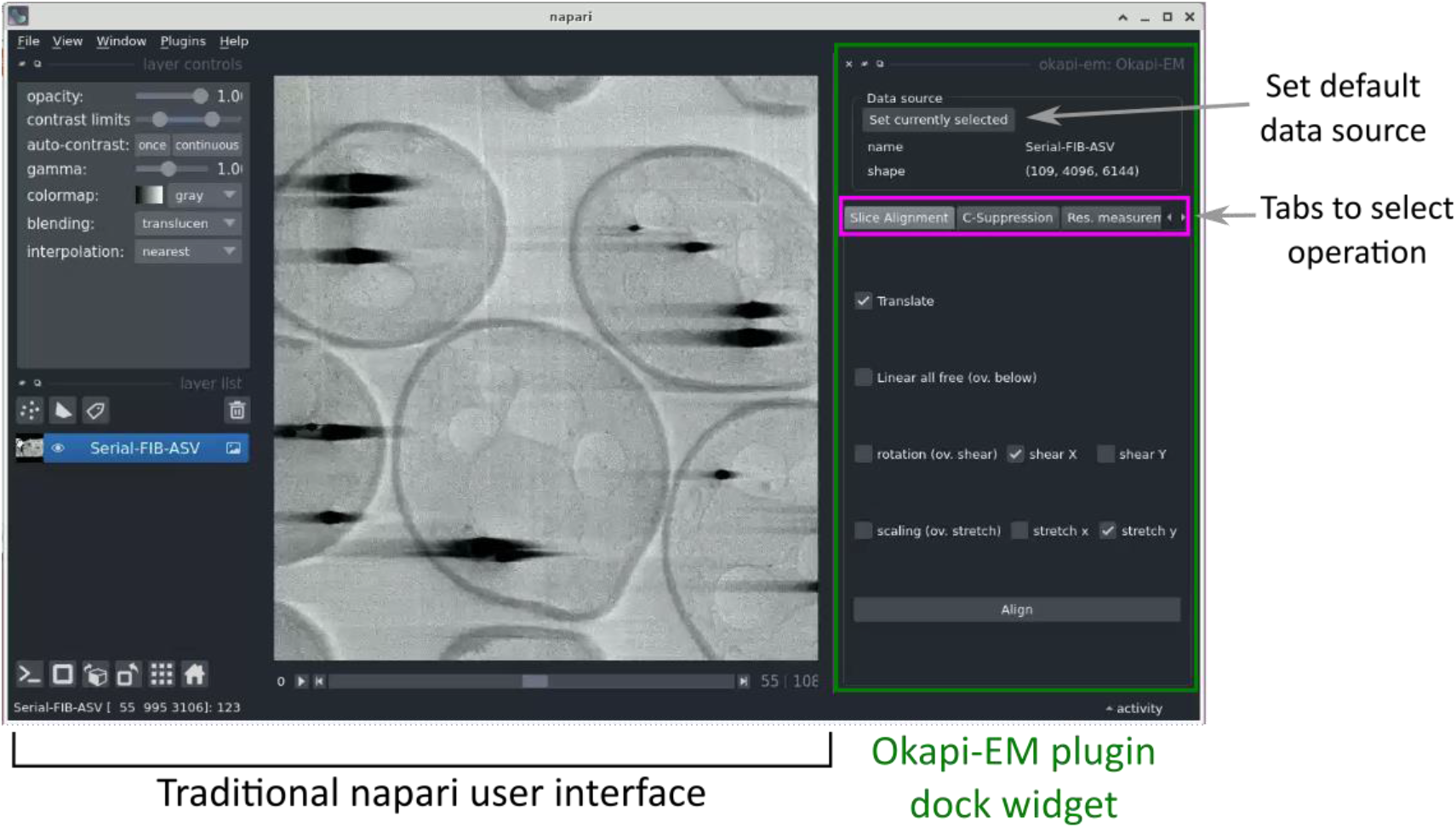
View of the Napari application highlighting the Okapi-EM plugin (green rectangle at right), and the currently available tools (pink rectangle at right). Each Okapi-EM plugin has its own tab with appropriate options for its use displayed. See Supplementary Figure 1 for a detailed view and description of the options available in each plugin.

### Stack alignment

In FIB-SEM, samples are milled in preparation for imaging. During this process, a number of issues can cause drift or misalignment in the image stack, such as the movement of the sample due to mechanical stress, small temperature variations and slight, iterative, stage movements going from milling to imaging positions that accumulate over time^(20)^. Inaccurate placement of the milling area by the operator or software may also lead to the observation of shearing during subsequent imaging^(21)^. These misalignments may further be amplified by factors including charging effects and instabilities caused by external disturbance^(21)^. As a result, an important element in the alignment of the resultant SEM images is the shearing or skewing along the direction perpendicular to the ion beam (or other layer removal option) trajectory^(20)^, which is particularly significant along the slow scan direction. Before any subsequent visualisation or analysis tasks, aligning the image stack is a crucial step.

While there have been efforts at improving image acquisition hardware and developing real-time correction^(22–24)^, these methods are limited by the need to ensure that the sample is not overexposed during data collection. Therefore, they do not provide fine alignment correction and require the use of subsequent alignment software^(21)^. Several software packages/plugins are available for this task, such as the closed-source and pay-for-service Amira by ThermoFisher^(25)^ and open-source options such as ImageJ plugins Linear Stack Alignment with SIFT^(26)^ and TrakEM2^(27)^. Although these software can in general align the images so that there is smooth transition between the slices, a major drawback is that they do not consider the physical process by which the images were acquired, thus can introduce distortions that are not present in the sample. During alignment, if incorrect transformation parameters are chosen, because slices are aligned to their neighbours, this can cause a cascade of transformations that can ultimately distort the shape of the stack^(28)^. Notably, the ImageJ plugins Linear Stack Alignment with SIFT and TrakEM2 do not offer options to perform alignment with shearing instead of rotation in particular directions, which is what images obtained by scanning line-by-line need to be compensated for.

### Alignment method

We have developed an improved image alignment method that aims to provide users with a variety of transformation options to appropriately describe image acquisition processes and to reduce the likelihood of introducing non-physical distortions during alignment of 3D stacks. It employs scale-invariant feature transform, or SIFT, a widely-used algorithm for the detection of local landmarks^(29)^. Since SIFT tends to include some false matches that could affect the alignment outcome, filtering is necessary to remove these outliers. Users can choose between percentile and RANSAC filters to achieve this. The former option is faster but less robust, while the latter is more computationally costly, but more reliable in cases where matches are scarce, or exhibiting misalignment. Users will then define a transformation that would best describe the physical distortion of the images depending on the image acquisition technique (Table 1) by adding or removing components of the transformation matrix, including translation, rotation, shearing, uniform scaling and stretching or shrinking, or selecting affine transformation which allows all six degrees of freedom. A description of each option is shown in Supplementary Figure S2. For instance, if the option shearing along the *y*-axis is enabled, and no scaling or shearing in *x*-axis are chosen, then the transformation matrix to be obtained is:

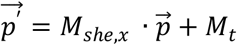

or:

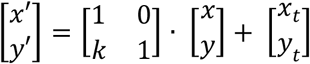

where 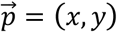 is a feature point on the source image and 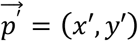 is its location after shearing with *M_she,x_* being the linear transform for shearing along *x* by the parameter of *k*, and translation of *M_t_* = (*x_t_, y_t_*). The distance between the 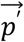 and the corresponding feature point on the reference image will be calculated. Using least square optimisation, the parameters *k, x_t_, y_t_* that minimises the summed distance will be found.

**Table 1:** Types of transformations that are commonly used to correct distortions from different cryogenic volumetric electron microscopy techniques.

|  | Translation | Rotation | Shear | Scale | Stretch | Affine |
| --- | --- | --- | --- | --- | --- | --- |
| **Cryogenic block-face imaging, laser sectioning (e.g., FIB/SEM)** | Yes | No | In scanning direction | No | In scanning direction | No |
| **Cryogenic block-face imaging, mechanical sectioning (e.g., SBF SEM)** | Yes | No | In scanning and sectioning directions | No | In scanning direction | No |
| **Cryogenic imaging of thin serial sections (e.g., ssTEM, array tomography)** | Yes | Yes | In sectioning direction | No | No | Yes |

### Rotational distortion introduced with existing software

As discussed previously, limiting the modes of transformation according to the physical process of image acquisition is crucial. Failing to do so when aligning FIB-SEM image stacks could result in a large angle rotation in the image slices that is not actually present in the sample. To demonstrate this, an artificial shearing transformation was applied to the original image (Figure 2A) acquired using cryogenic serial pFIB-SEM to mimic a shear force distortion (Figure 2B). For this image pair, using SIFT, many matches can be found on the left side of the image, especially in the triangular and circular region (Figure 2C). Using least square or other optimisation methods without restraining the transformation mode results in a large angle rotation or an affine transformation that aligns those feature-rich regions well, but fails to align other parts of the image and introduces a non-physical distortion (Figure 2C right).

**Figure 2.**
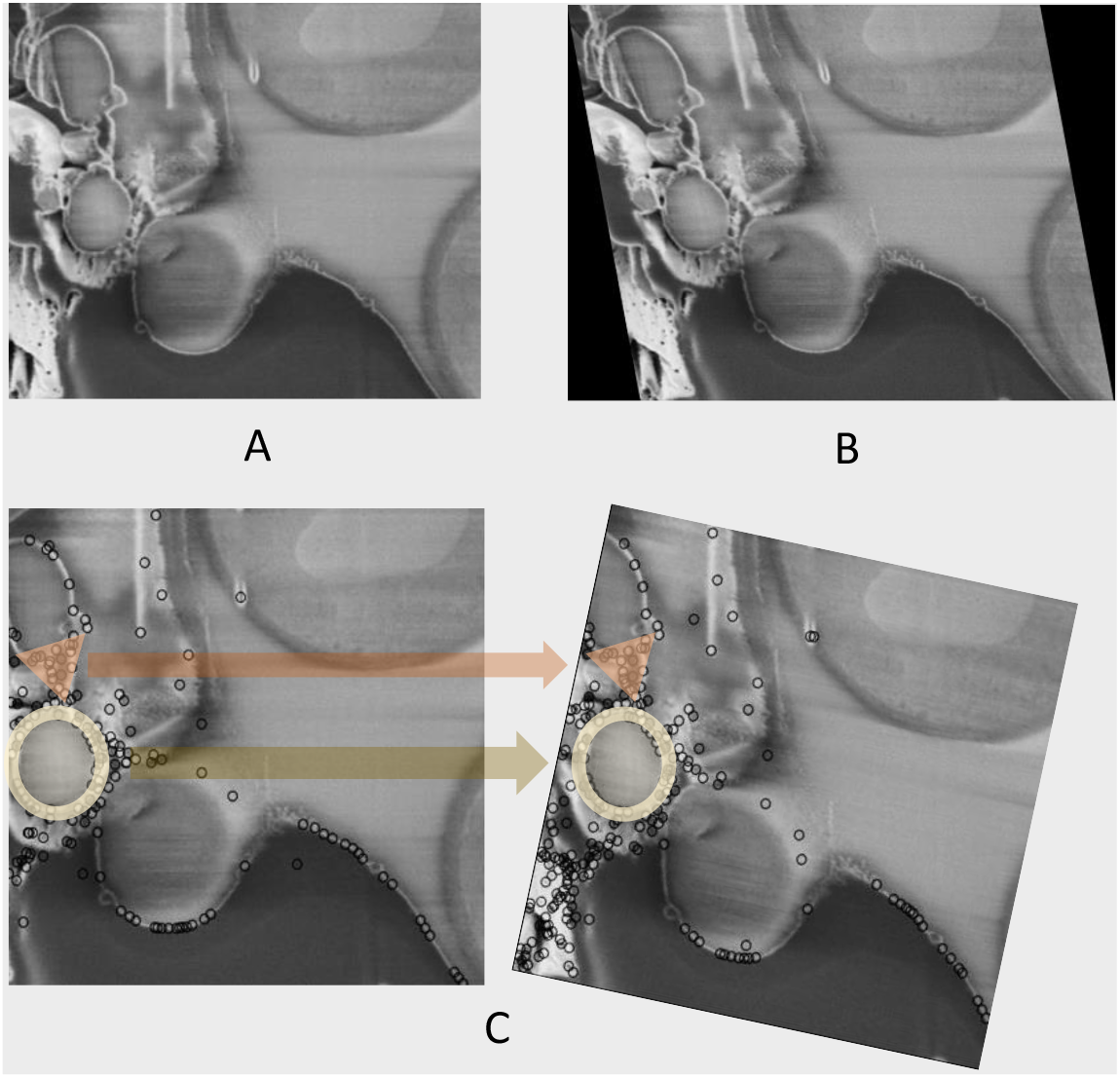
Non-physical distortion in the alignment process if the modes of transformation are not appropriately limited. (A) Original SEM image of yeast cells. (B) Artificially distorted image after a shearing transformation was applied to the original image. A shear factor of 0.15 was selected for visibility. (C) Feature points found in (B) using SIFT, with two feature dense regions highlighted with circle and triangle markers. (D) Possible alignment result when the modes of transformation are not restrained, where even though the feature-rich regions are well-aligned, a non-physical rotation is introduced. Data is EMPIAR XXXX.

## Results

Okapi-EM stack alignment was applied to a cryogenic serial pFIB/SEM image stack of yeast cells (109 slices). As described in Table 1, translation, shear x axis and stretch along y axis were chosen to compute an alignment based on the landmarks found using SIFT with RANSAC outlier filtering. Cross-sectional views are used to visualise the alignment result (Figure 3). To compare outcomes, the same image stack was aligned in Fiji plugins TrakEM2 and Linear Stack Alignment with each transformation option offered as well. Substantial drift and distortion can be observed in the unaligned stack (Figure 3B, 3J; Supplementary Movie 1). After alignment, cell shapes are restored with mostly smooth outlines (Figure 3C, 3K; Supplementary Movie 2). While all alignment approaches improved the cross-sectional alignment to some degree, distortion can still be observed in the outline of the cells when using either TrackEM2 or Linear Stack Alignment (Figure 3 D-I, L-Q), especially at the y = 2 μm when using TrakEM2 and when affine transformation is selected in Fiji Linear Stack Alignment.

**Figure 3.**
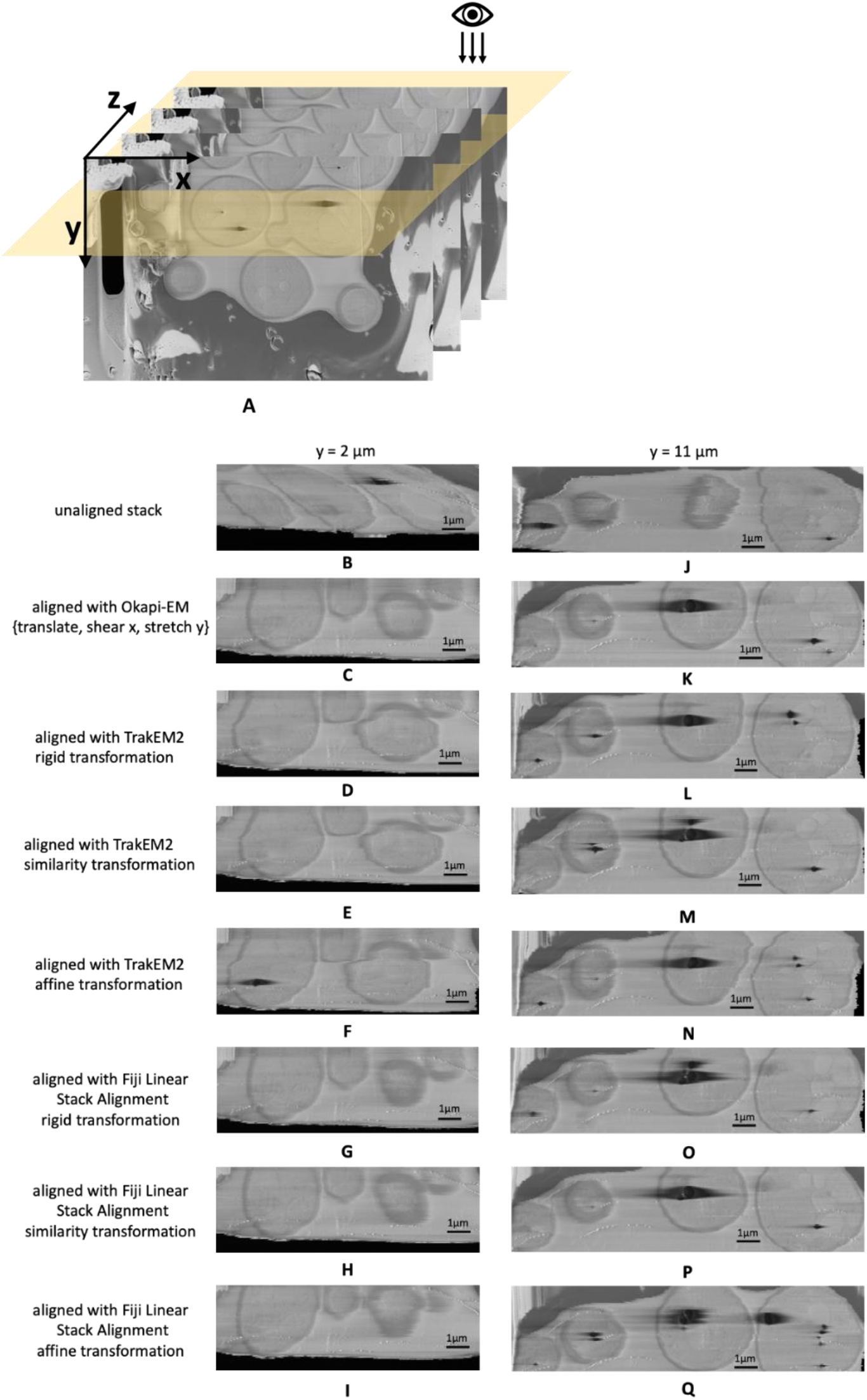
Alignment results and comparison between Okapi-EM, TrakEM2 and Fiji Linear Stack Alignment with SIFT (A) Cross-sectional views of SEM image stack of yeast cells (109 slices along z direction, slice size 20.7×13.8 μm^2^) are obtained. (B)-(I) Cross sectional view of a 11.2×3.7 μm^2^ area at y = 2 μm of the unaligned stack and the aligned stacks using Okapi-EM alignment with RANSAC and {translate, shear x, stretch y} selected, Fiji plugin TrakEM2 with rigid transformation, similarity transformation, or affine transformation selected, respectively, and Fiji plugin Linear Stack Alignment with rigid transformation, similarity transformation, or affine transformation selected respectively. (J)-(Q), Cross sectional view of a 15.0×3.7 μm^2^ area at y = 11 μm of the aligned stacks using the three aforementioned alignment methods and settings respectively. Data are EMPIAR XXXX. In both Fiji plugins, rigid transformation allows translation and rotation. Similarity transformation allows translation, rotation and scaling. Affine transformation is defined as shown in Supplementary Figure S2.

### Charge artifact mitigation

Charging artefacts often appear when insulating materials interact with the electron beam^(2,4)^. This effect is normally minimized by adjusting beam energy or scanning parameters, and its severity depends on the target substance being imaged. Biological samples can present a challenge, with cells often containing different compartments with distinct electrical conduction properties that cannot be completely balanced through adjustments in acquisition parameters. Inevitably, images will show evidence of charging, causing artefacts which manifest in the SEM images collected in the form of elongated dark regions in the scanning direction that extend asymmetrically beyond the charging feature itself (Figure 4A and 4G). The practical outcome of this artefact is the partial obscuring of biological features nearby to insulating materials. This makes both manual and automated downstream data processing and analysis (e.g., segmentation or visualisation) more difficult.

**Figure 4.**
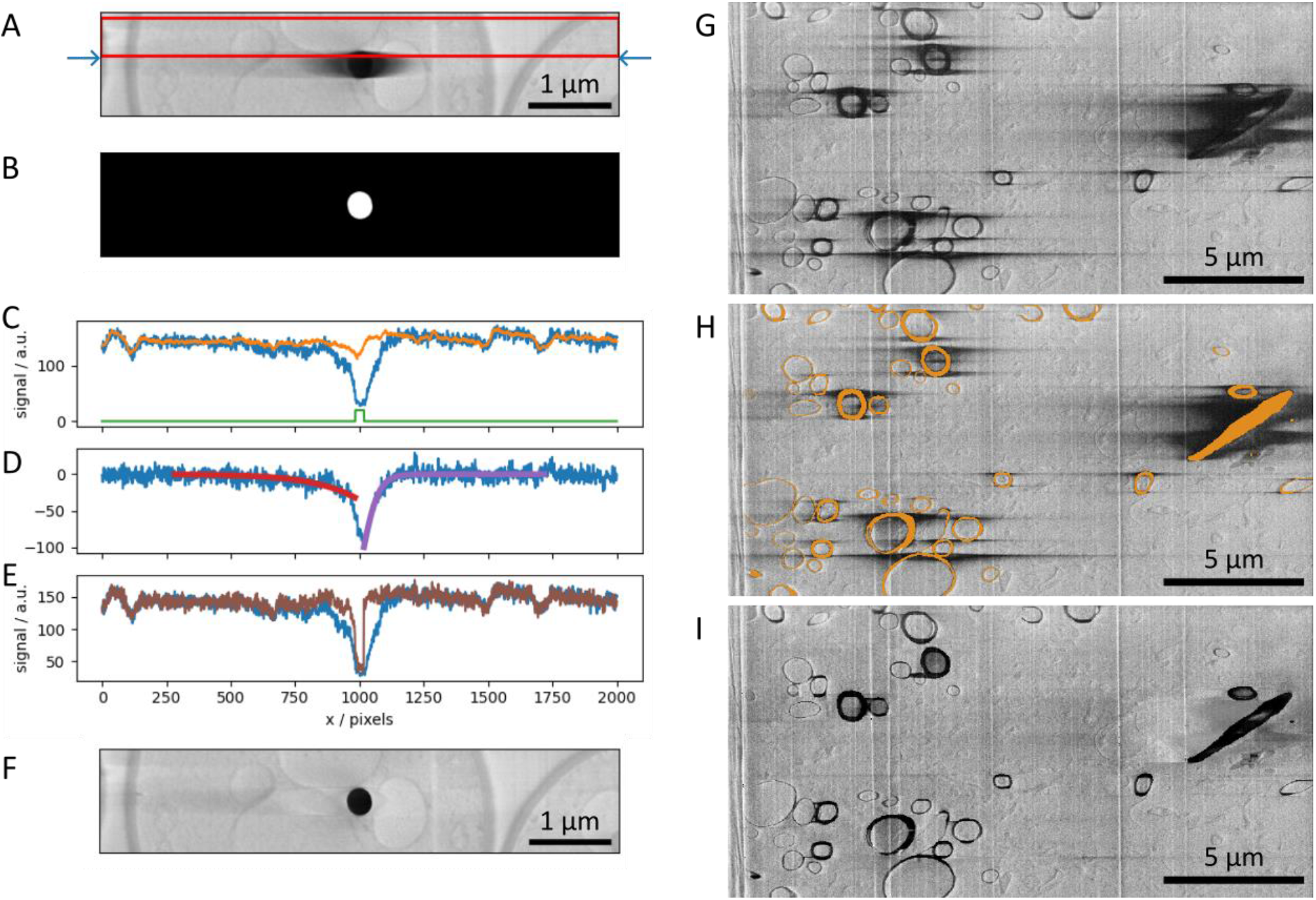
A-F SEM images and plots illustrating the charge artifact removal algorithm on a lipid droplet from a yeast dataset. Data are EMPIAR XXXX. (A) SEM image of a lipid droplet within a yeast cell (image size 6.75×1.35 μm^2^). Arrows indicate a scanning line of interest, with its signal profile in (C-E) as blue line. Red rectangle represents data region where signal was averaged resulting in signal profile in (C) as orange line. (B) Annotation image of the charge centre, corresponding to the charging artifact in Figure A. Green line in plot (C) is the line profile of the annotation along the row of interest (arbitrary units). (D) Line profiles describing the background corrected signal (blue line) and the optimised functions f_left_ and f_right_, (red and purple) respectively. (E) Line profiles of the uncorrected signal (blue line) and corrected signal (brown line). (G-I) SEM images illustrating charge artifact suppression on myelinated sheaths found in mouse brain (image size 20.7×10.3 μm^2^). Data are EMPIAR XXXX. (G) Original SEM image displaying copious amounts of charging. (H) SEM image with overlaid segmentation of charging centres and extending to complete rings of myelin sheaths, (I) result after applying filter with default parameters.

A previous attempt to mitigate charging in SEM images^(4)^ utilises the python scikit-image dilation/morphology function, based in an algorithm proposed by Robinson and Whelan^(30)^. This method separates the charging tail signals from the SEM images and then partially subtracts them from the original image. In our hands, this method did not work with the charging artifact tails in our datasets (Supplementary Figure S5), possibly indicating it is specific to the datasets it was developed for or the instrument used to collect them.

Okapi-EM contains a charging artefact suppression tool (chafer^(31)^) which is designed to restore the image contrast within and around charging artefacts, while retaining the charge centres themselves. As the artefact appears elongated along the direction of scanning (laterally) and is therefore influenced by the rastering nature of the scanning, it immediately suggests that to reverse this effect in the images, a row-by-row filter that uses information of the surrounding areas, in particular along the direction perpendicular to scanning, should be used to subtract the charging effect. With Okapi-EM’s chafer, the restoration method operates sequentially row-by-row in down and up passes, and the estimation of the charging artifact tails is done by fitting with a smoothing function, which is later subtracted.

For this filter to work a semantic segmentation of the charging centres must be provided. This segmentation can be done manually or by using shallow or deep-learning ^(32)^ predictors using either tools within Napari or elsewhere^(25,33)^. In our experience, simple thresholding methods to label either charging centres or full artifacts don’t work well due to the presence of other features of similar intensity. Correct annotation of the charging centres is crucial as it provides both an inverted mask of charging locations for correction and an indication of where the charging artifact is relative to the charging centre, hence being useful for choosing initial values during optimization of the functions used (see below).

The filtering scheme works in the following way: First it takes the previous rows (Figure 4A, red box) and averages perpendicular to the scanning direction (vertical here). This is used as an approximation of the signal without the charging effect. The difference between the current row (Figure 4A, blue arrows) and the previous-row average is assumed to be an estimation of the effect of the charging (Figure 4C-E, blue line). Simply subtracting this estimation of charging effects from the current row of data gives noisy results from row-to-row. Instead, the tails of the charging effect signal (artefact that is outside of the charging centre) can smoothed or modelled by curves, and when subtracting this from the original signal, it gives results that are more consistent from scanning row to row.

We have trialled several functions that may fit and optimize best to the charging artifact tail signals. Exponential (*Ae^−kx^*) and gaussian functions 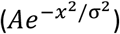 can fit reasonably well to these tails but overall give poor results when reconstructing images, in particular at the function optimization step the gaussian filter often fail to converge to suitable parameters (see Supplementary Information Figure S4). Some of the reasons why it fails to converge is that the signal is often flat in the rows above the charging artifacts, while other times, there are other biological features nearby that interfere with the function fitting, leading to unrealistic curve fitting parameters. Instead, we found that a shifted logistic sigmoid function (or Fermi-Dirac distribution type) works very well for this task. This function appears like a smoothed step function and is widely used in machine learning algorithms as an activation function, having the advantage that it ‘saturates’ on either side of the curve, which is more characteristic of the charging artifact tails observed. Because the tails themselves were asymmetrical, the functions used to mitigate them were different for the left side and right side of the charging artifacts, and given by:

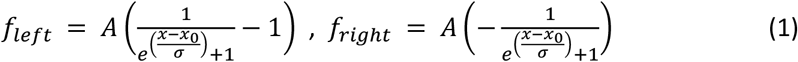

with *A*, *x*_0_ and *σ* being parameters to be fitted, and functions *f_left_* fitted on the left of the labelled artefact, and *f_right_* fitted on the right side.

After optimising *f_left_* and *f_right_* (Figure 4D, red and purple curves respectively) these functions are subtracted from the row signal (excluding masked regions), resulting in a charge mitigated signal that still represents the charging object (Figure 4E, brown). Running this process row-by-row in up-down and down-up passes results in a filtered image (Figure 4F), which is a significant improvement compared to Figure 4A, suppressing the main artefacts, while recovering some of the washed-out data previously hidden below.

In addition to SEM images of lipid droplets found in yeast cells, a second dataset featuring myelinated sheaths from mouse brains was used to test this process (Figures 4G-I). The manual segmentation of charging centres (Figure 4H) included both the charging centres and whole organelles, despite many regions not displaying charging artifact tails (including these non-charging regions has no negative impact, see Supplementary Information Figure S6). The filtered image (Figure 4I) demonstrates a substantial visual improvement to the image quality. We note that in regions near significant charging effects, although the contrast can be matched to the remaining image, there are no recoverable biological features present. This is particularly noticeable around the elongated diagonal organelle on the right side of the image.

The main advantage of mitigating charging artifacts mitigation is the improved contrast of organelles near the charge centres. As such, both manual and automatic segmentation tools are expected to perform better when charging artifacts are absent, as these are notoriously sensitive to contrast changes which could lead to poor visualisation and missing or misidentified organelles.

To better understand the effects of our charge mitigation scheme, we have compared the background in the vicinity of charging centres in corrected images (e.g., 4F or 4I) to their uncorrected counterparts (e.g., 4A or 4G), taking care to exclude regions that have been labelled as charging centres from the calculation. Through filtering, we expect to “uncover” biological features that were previously obscured by the charging tails and as such would expect a decrease in the standard deviation of the signal intensity around the charging centre. The standard deviation of the signal intensity of the results shown in Figure 4F is 60% lower than the signal intensity in Figure 4A. Similarly, the standard deviation signal in Figure 4I (excluding labelled regions) is 58% less than in Figure 4G, suggesting the regions previously obscured by the charging artefacts are now in a more comparable range of signal intensity to the regions throughout the rest of the dataset. Through visual inspection, it is clear that while some information is recovered, at close proximity to the charging centres information is lost due to the presence of the charging artefact.

### Resolution estimation using one-image FRC

Measuring image quality enables users to evaluate the best imaging protocols for the requirements of their research question. Image resolution is a helpful measure of image quality as it can be directly related to physical dimensions, so it is intuitive to interpret. Resolution can be defined as “a maximum spatial frequency at which the information content can be considered reliable”^(34)^.

One method of determining image resolution is via a two-image Fourier Ring Correlation (FRC)^(35)^, which has been implemented in fluorescence microscopy, and is modified from single particle methods in structural biology. It is both a measure of resolution and consistency between images within a dataset. In this method, two independently acquired images of the same field-of-view are compared in the frequency domain to find the highest frequency where the images can be said to be similar. This is performed by determining the frequency where cross-correlation drops below a given threshold, often 0.143^(36)^. The requirement for two independently acquired images is challenging to fulfil as it cannot be done retrospectively, and repeated acquisition could introduce image artefacts (i.e. beam damage), affecting the resolution.

Koho and colleagues proposed a one-image FRC calculation based on sub-sampling a single image to produce pairs of images^(37)^. This method arranges alternate pixels of an image in a checkerboard pattern to create sub-images from a subset of pixels and then calculates the FRC between the sub-images. A calibration function is then applied to match the one-image FRC from sub-images to the gold-standard two-image FRC for the specific microscope and imaging conditions used.

The method of Koho *et al*. has been adapted for serial FIB/SEM imaging in a software tool called Quoll^(38)^. Quoll is an open-source, user-friendly tool, and library to calculate the local resolution of single images with the one-image FRC. This method can be applied to any imaging modality once the calibration has been performed. The 2D image is first split into tiles, and the one-image FRC is calculated on each, returning a map of the spatial variation of resolution across the image, and a plot of its distribution. This process can be done on multiple 2D images within a 3D stack to understand resolution throughout the dataset. If large regions of featureless background or artefacts are present, it could skew the output as measurements would be taken on areas without appropriate levels of information present; these can be excluded through masking prior to assessment.

### Calibration

Calibration is required to match the one-image FRC to the gold-standard two-image FRC resolution. This is an instrument-specific calibration as it is affected by the noise generation of the instrument. The calibration dataset requires two repeated images of the same field of view taken at several pixel sizes/magnifications. Ideally, there should be no significant, artificial changes between the images such as artefacts or image shifts, as these could affect the two-image FRC that is calculated between them. If necessary, image registration can be used to correct shifts and choosing specimens which do not suffer from specimen degradation through repeated imaging is helpful (i.e., inorganic options such as gold or polystyrene beads).

In our recent cryogenic pFIB/SEM study^(2)^, calibration was performed using images of polystyrene beads of 1 μm diameter (Abvigen, USA). The images show the same field-of-view at 1.12, 2.25, and 4.5 nm pixel size. The final dataset consisted of 6 images, with a pair of images taken at each pixel size and SEM angle. The linear stack alignment with SIFT plugin in Fiji 2.3.0/1.53q was used to register each pair of images to each other, to ensure the exact same field-of-view was considered for both images^(39)^. The images were cropped to cover the same physical field-of-view for all pixel sizes, 2000 x 2000 nm for the 52° images and 1600 x 1600 nm for the 90° images (Figure 5). The number of pixels in each pair of images was different due to the varying pixel size.

**Figure 5.**
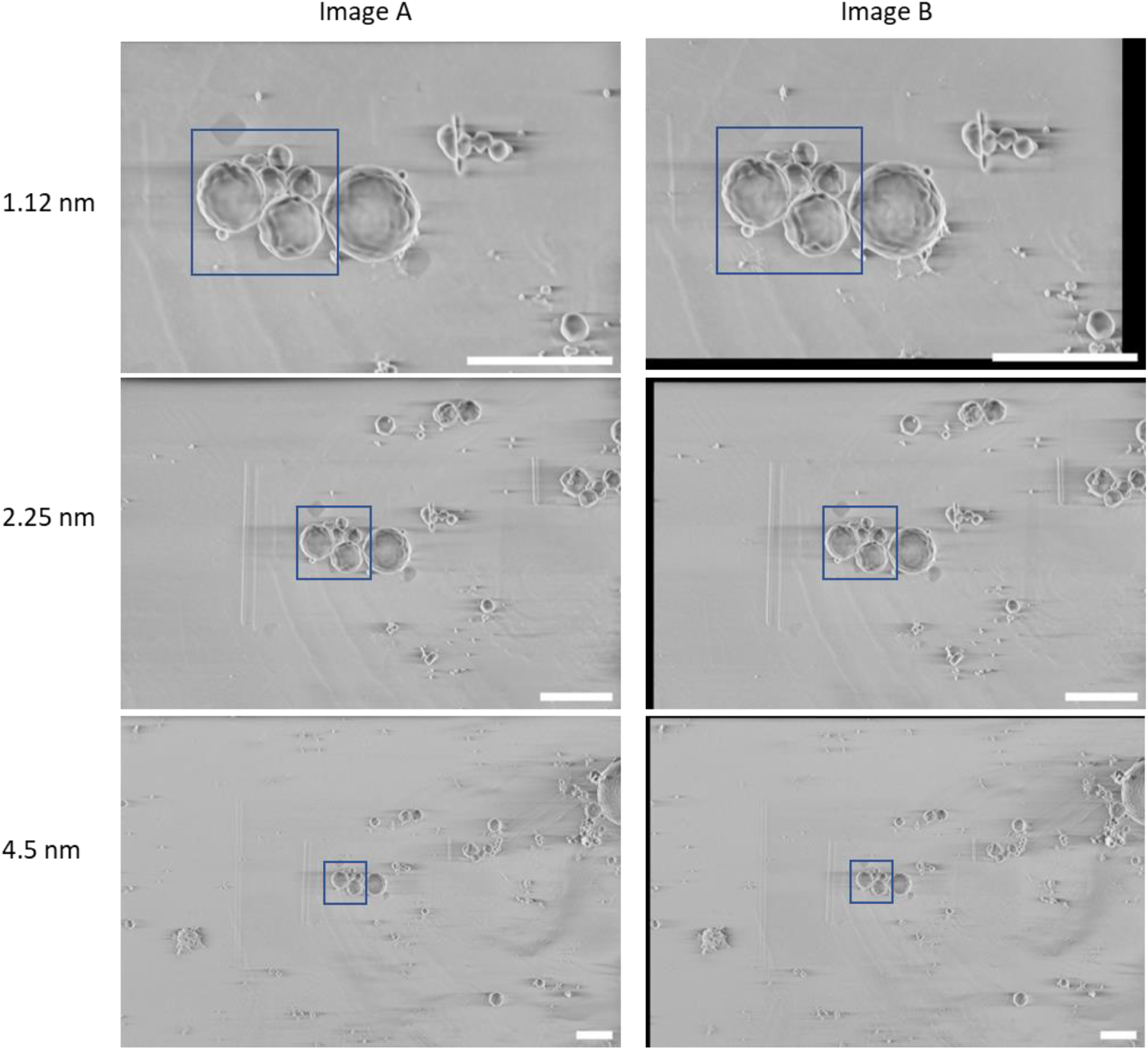
Calibration dataset of polystyrene beads used to calculate the gold-standard two-image FRC. These images were taken at 52° SEM angle. A region-of-interest (blue box) covering 2 μm x 2 μm was used for the calibration, where the average one-image FRC and two-image FRC were calculated for these regions at all pixel sizes. Scale bars represent 2 μm.

The FRC curve (cross-correlation vs. frequency) was calculated from each pair of images, and normalised to a scale of 0-1, where 1 was the maximum frequency calculated. This normalisation enabled direct comparison of FRC curves between images. The normalised frequency at which the cross-correlation fell below 0.143^(36)^ was taken as the reference resolution. The one-image FRC resolution was also calculated following the checkerboard sampling method of Koho and colleagues, this was the *r*_*co*1_.

A calibration function was fitted to the plot of 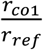 against *r*_*co*1_ (Figure 6). This calibration function shifted the one-image FRC curve to match the two-image FRC, so that the resolution from both methods were comparable. This calibration function is instrument-specific, so it can be reused for any images taken from that instrument, provided that the image acquisition parameters are within the range of the calibration dataset.

**Figure 6.**
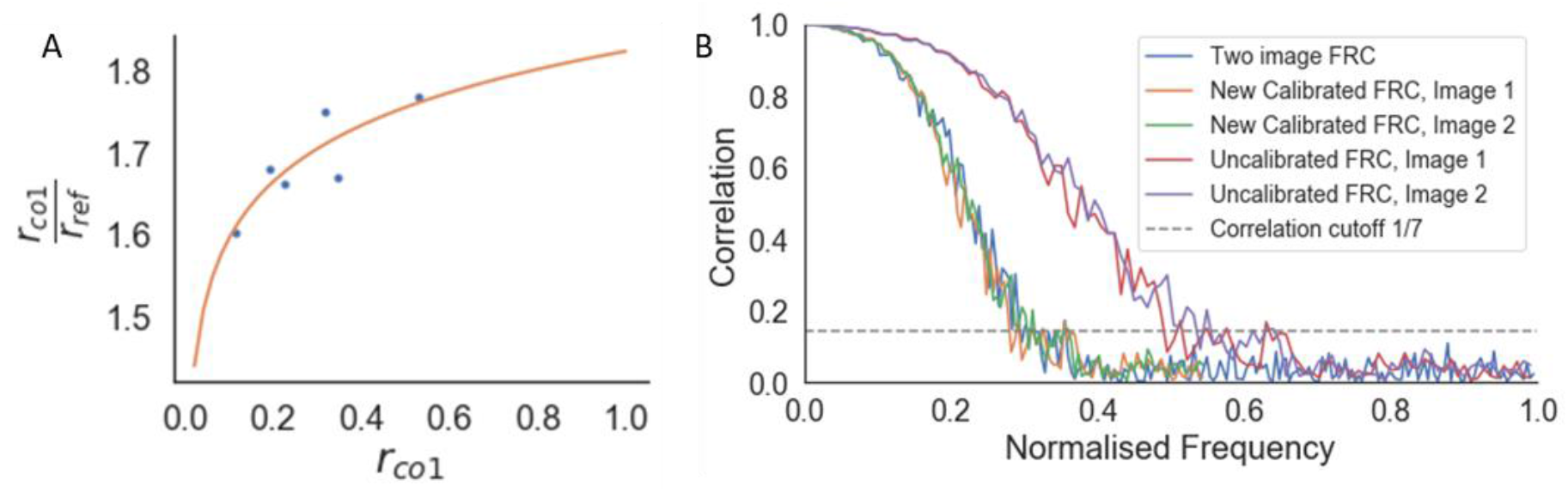
Calibration of the one-image FRC measurement to the gold-standard two-image FRC. (A) Calibration curve 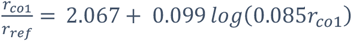, where r_co1_ is the average resolution measured from the one-image FRC for the image pair, and r_ref_ is the resolution from the gold-standard two-image FRC measured from the image pair. (B) Two-image FRC curve and one-image FRC curves before and after calibration for the image pair at 4.5 nm pixel size at 52 ° SEM angle. Calibration shifted the uncalibrated one-image FRC curves along the frequency axis to match the two-image FRC curves, ensuring that the resolution measurement for the one-image FRC matches the gold-standard.

### Application to cryogenic serial FIB/SEM data

Quoll was used in a recent publication to develop cryogenic serial pFIB/SEM imaging of biological specimens^(2)^. Here, Quoll resolution measurement was demonstrated on five different biological specimens (*R. rubrum*, HeLa cells, Vero cells, mouse brain, *S. cerevisiae*). The measurements were used to show that imaging at 90° SEM angle produced better results than 52°, to determine the depth of field of the instrument, and to show that there was no degradation of image quality through serial sectioning of the specimen, even at depths of ≈25 μm. These performance evaluations would not have been possible without the quantitative measurements from image resolution estimation with Quoll.

The image resolution reported by Quoll was validated against biological structures with known physical sizes. The local image resolution was calculated for a cryogenic serial FIB/SEM image of HeLa cells on tiles measuring 256 x 256 pixels, which corresponded to a physical field-of-view of 1.73 μm x 1.73 μm. These images contained nuclear pore complexes and centrosomes, which are approximately 120 nm and 200 nm in diameter respectively^(40,41)^. These structures were clearly resolvable in the images, and resolution in the tiles containing these structures surpassed the known diameters of these structures, validating the resolution measurements (Figure 7). FRC measurements are carried out on tiles within image slices and the reported measures provide a plot of the mean and standard deviation of resolution across the slice, as well as a heatmap for visualisation. It is important to emphasize that these values are not the highest resolution visible within the slice, but instead a representation of the mean resolution of each tile. The goal of this method is to provide a quality metric for the raw data as a whole.

**Figure 7.**
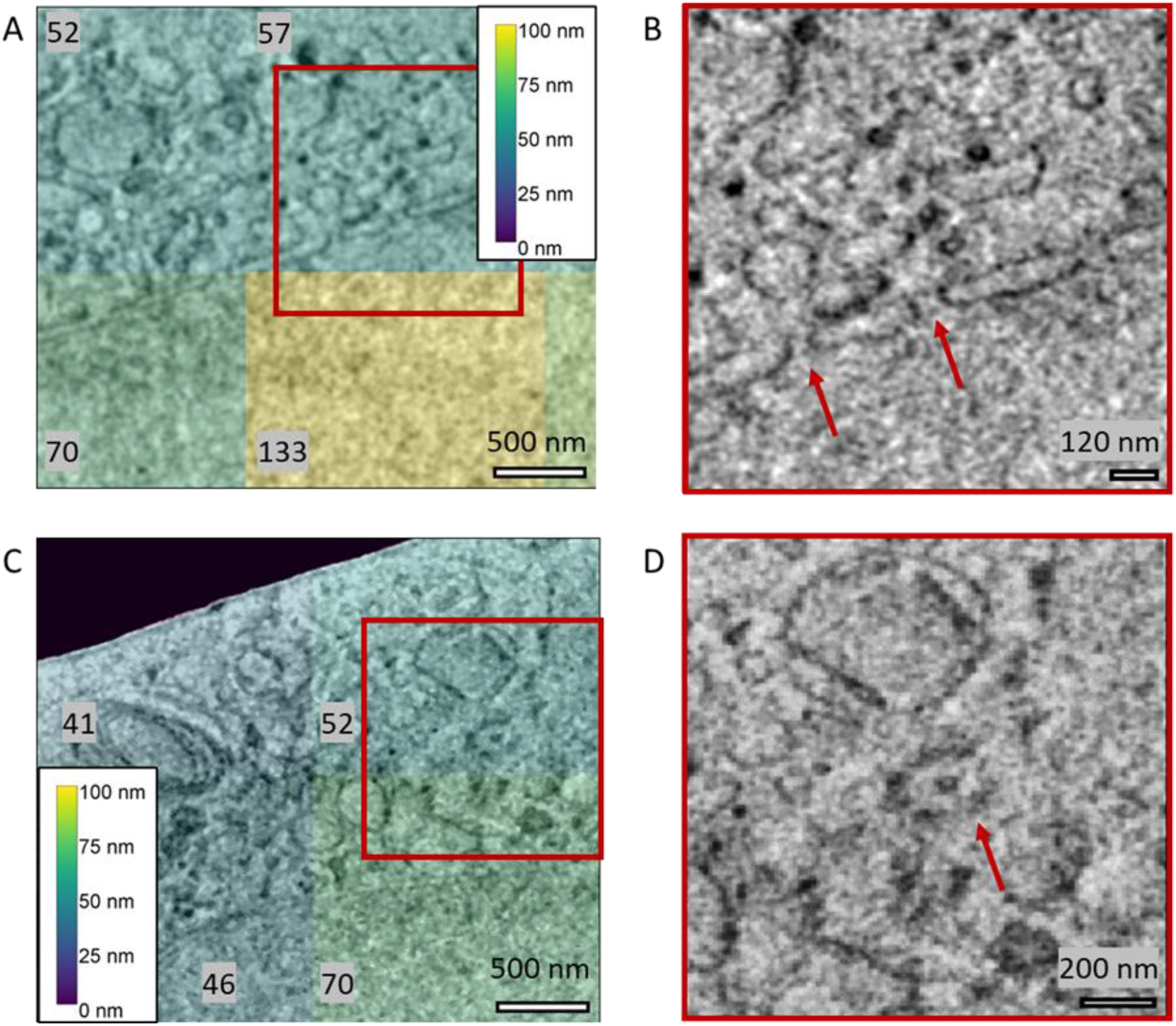
Validation of Quoll resolution measurements compared to physical features of known sizes in HeLa cells. Nuclear pore complexes (A, B) of 120 nm diameter were clearly resolved in the images, as indicated by the arrows in (B), and the resolution in this region was estimated as better than the resolvable features. Similarly, in (C) and (D), centrosomes of 200 nm diameter could be resolved from the images (indicated by the arrow in D), and resolution was better than the size of the centrosomes. (A, C) A resolution heatmap is overlaid onto regions of the image showing nuclear pore complexes and centrosomes respectively, where the colours represent the resolution values of that local region. (B, D) are the zoomed-in regions indicated in (A) and (C) with a red square, with arrows indicating the position of the nuclear pore complexes and centrosomes respectively. Data is EMPIAR XXXX.

We found that tile size does not affect the overall resolution distribution of the image. The local image resolution was calculated on cryogenic FIB/SEM images of *S. cerevisiae* and *R. rubrum*, at 3.37 nm and 1.94 nm pixel sizes respectively. The images were sampled with tile sizes of 128 × 128, 256 × 256, and 512 × 512 pixels. The difference in minimum and maximum median resolutions for all tile sizes was within 0.38—0.48 pixels. The Kruskal-Wallis H-test was applied with the null hypothesis that the population median of all groups was equal^(42,43)^. The null hypothesis could not be rejected (p > 0.05) so the median resolution for all three tile sizes was considered equal.

## Discussion

Cryogenic serial FIB/SEM imaging provides exciting opportunities for in situ structural biology, though as a relatively new method of imaging, requires development of appropriate computational tools both for assessment of the method and processing of the data to enable biologically relevant qualitative and quantitative outcomes. Okapi-EM has been developed to begin this process. It includes three plugins to align serial SEM stacks, mitigate charging artefacts and to assess the resolution of SEM data. In future, Okapi-EM will also include a separate plugin to measure and mitigate curtaining artefacts.

Many of the approaches developed here for cryogenic serial pFIB/SEM are likely also applicable to room temperature FIB/SEM, non-plasma-based serial SEM (SBF-SEM) or serial TEM (serial section TEM or array tomography) with little or no modifications needed to the method or implementation. If modifications are needed, we are happy to adapt Okapi-EM to meet these needs and encourage feedback from users and developers through contacting the corresponding author here or via our GitHub page.

Okapi-EM will continue to be developed into a more automated, quantitative workflow for data processing in the future. For serial FIB/SEM, alignment of 3D stacks is generally one of the first data processing steps and its outcome can have an impact on all downstream processing and analysis. It is therefore important to apply the minimum amount of transformations (i.e., change the data as little as possible), but do enough to ensure the data is interpretable. Finding this balance is currently a manual process. Development of image alignment quality metrics will enable this to shift to an automated process in the future.

Currently, the next step in data processing for serial FIB/SEM data is mitigation of artefacts such as charging. As described above, our implementation relies on the prior identification of charging centres through either manual or machine learning-based segmentation approaches. Our focus in this area will be on the development of a more automated approach to charge centre identification and segmentation. Additionally, we would like to extend our charge mitigation approach to data which display charging in visually different ways. We are aware of charging which is characterised by artefact tails that are bright white surrounding black charging centres or a mixture of white and black charging artefact tails again surrounding black charging centres. Further testing of the filters suggested here, and others is necessary to identify or adapt mitigation strategies for these data types. However, this testing has been hindered by lack of access to publicly archived example data (e.g., in EMPIAR).

Finally, once the data has been aligned and artefacts mitigated, the next steps are visualisation, assessment and segmentation. The FRC resolution measurement is a first step towards quantitative assessment of the output data though others related to “segmentability” (i.e., contrast, information content, presence/absence of a feature of interest) could be developed.

It would also be valid to use image quality assessments at the beginning of the process – data collection - guiding users in optimising their imaging protocols, where acquisition settings can be determined based on the specified quality metrics and the researchers specific question(s). This is especially helpful for new users who may not have the experience to quickly determine the optimum combination of acquisition settings, so image resolution and other quality measurements at the microscope could guide the user to quantitatively optimise their protocols, removing some of the superstitious/anecdotal approaches that are sometimes found in research.

This thought process can then be extended to automated microscopy methods, where acquisition settings are adjusted during serial imaging to obtain good quality images throughout the session even as the sample features change. This would enable longer image acquisition sessions as the user is not required to manually adjust the imaging settings throughout the session. Real-time measurements and parallelised computational approaches will be required for these assessments to quickly identify and react to issues during imaging.

Image quality information can also be saved as metadata with each acquired image or stack of images, which is useful for indexing the images for future re-use. For example, future users could search for images of specific specimen types with a minimum resolution to answer their research question, or methods developers could search for images with low resolution to explore how their methods could improve existing images.

## Conclusion

Volume EM (room temperature or cryogenic) provides exciting opportunities for visualisation and quantification of cellular and tissue components, though in many cases, the outcomes of these studies are hampered by the manual nature of data processing and analysis. We have provided a bundle of plugins within the Napari data visualisation package to speed the process and ease the burden on researchers using serial SEM or TEM imaging techniques. These tools allow for the alignment of 3D stacks without introduction of unnecessary transformations, the mitigation of charging artefacts caused during scanning imaging techniques and the assessment of data quality through one-image resolution measurement. These tools are in regular use and development, and we welcome feedback and contributions as they are extended to new data types, imaging modalities and purposes.

**Table 2:** Summary statistics of resolution distribution measured from different tile sizes. Tile size was not found to affect the overall resolution distribution of the image. The resolution distribution median was found to be equal for all tile sizes (p > 0.05) by the Kruskal-Wallis H-test. All measurements rounded to 3 significant figures.

| Tile size | Number of tiles | Mean Resolution (nm) | Standard Deviation of Resolution (nm) | Median Resolution (nm) |
| --- | --- | --- | --- | --- |
| ***S. cerevisiae*, pixel size 3.3724 nm** |  |  |  |  |
| **128** | 256 | 45.2 | 9.68 | 45.3 |
| **256** | 64 | 47.5 | 10.4 | 46.6 |
| **512** | 16 | 44.0 | 6.96 | 45.3 |
| ***R. rubrum*, pixel size 1.94171 nm** |  |  |  |  |
| **128** | 64 | 20.9 | 5.31 | 19.7 |
| **256** | 16 | 21.1 | 5.31 | 20.6 |
| **512** | 4 | 20.3 | 1.67 | 20.3 |

## Supporting information

serial-FIB/SEM of yeast - unaligned stack

serial-FIB/SEM of yeast - aligned stack

## Authorship contributions

### Manuscript

L.M.A.P, E.M.L.H and Z.C.C. wrote the manuscript in consultation with M.B, M.C.D. All authors reviewed the manuscript prior to submission.

### Software development

L.M.A.P. for Okapi-EM graphical user interface plus bundling and chafer filter, E.M.L.H. for Fourier ring correlation software (Quoll), Z.C.C. for SIFT stack alignment, N.B.-y.Y. for general discussions and ongoing development of additional tools.

### Data acquisition

M.D. for the yeast data, T.G. and M.G. for mouse brain data, L.W. for the FRC calibration dataset; Project management: M.B, M.C.D.

## Data Availability

Sample data used for alignment will be made available on EMPIAR alongside Dumoux et al.^(2)^.

## Supporting Information

**Figure S1.**
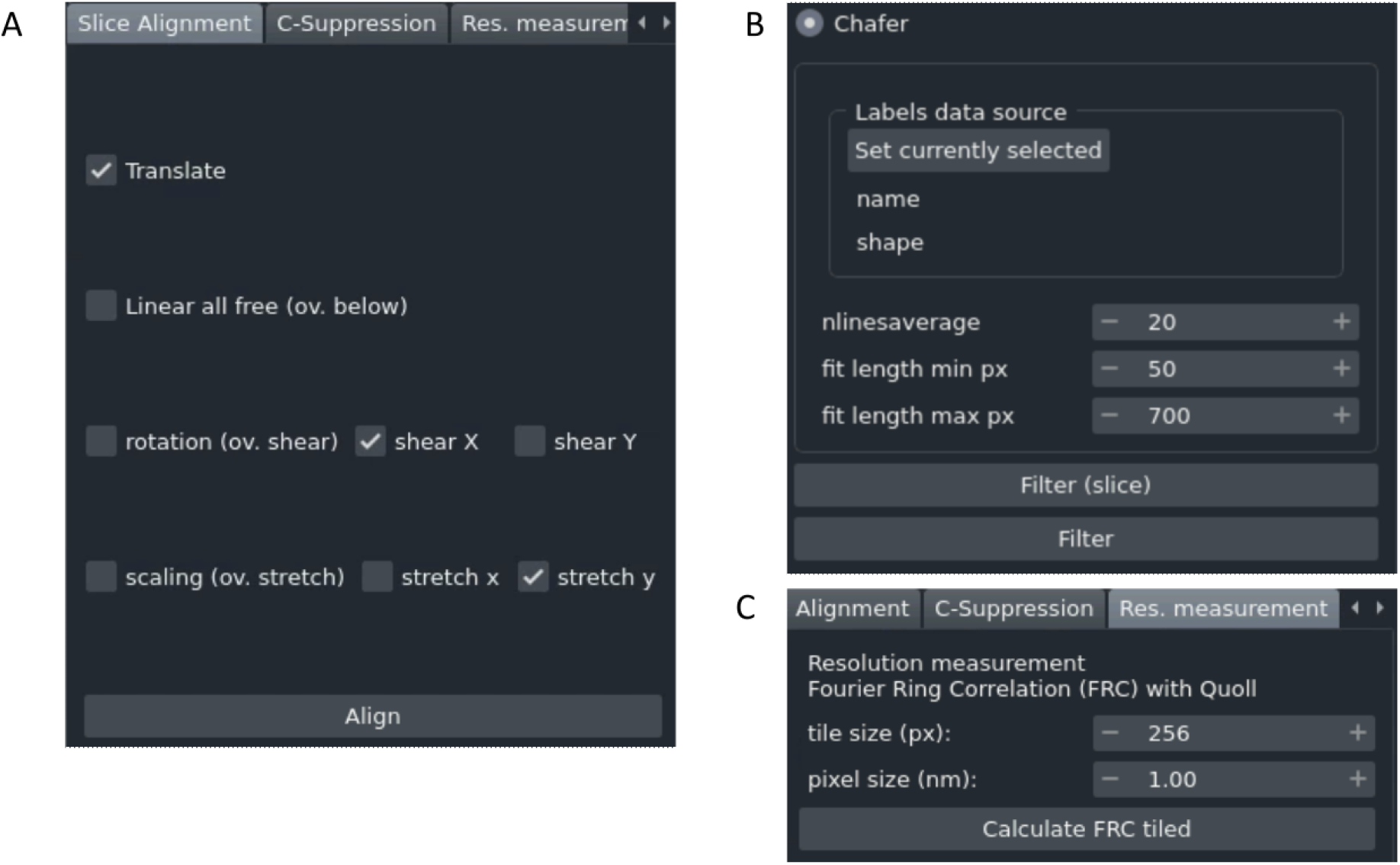
The graphical user interfaces for all three Okapi-EM plugins. (A) In the slice alignment user interface, the user can choose which parameters in the transformation(s) are allowed to be optimized. Defaults are freedom to translate, shear x, and stretch y, which are best for SEM slice stacks. (B) In the charge artifact removal filter interface (chafer) the user can select the labelled charge centres and specify parameters related to the filters. (C) In the resolution measurement interface (Quoll) the user must specify the tile size (in pixels) and the pixel size (in nanometers). After execution, a new napari layer will appear as a heat map overlay, and a statistical summary of the calculated values will appear in the user interface.

The user interface for the slice alignment tool (Figure S1A) offers several options that determine which and how the parameters in the transformation matrix between image slices are optimised. Linear all free (also known as affine in other software packages) means that all parameters in the linear transformation are adjustable during the optimization. If the rotation option is selected, rotation is optimized, but shearing along X or Y axis are not (overridden), being equivalent to *rigid body* in other alignment algorithms. If rotation is deselected, shearing along x or y directions can be individually set as adjustable. If scaling option is selected, a single scaling value for both horizontal and vertical directions is adjustable during optimization. With this option deselected, the scaling along vertical and horizontal directions can be allowed/disallowed individually.

The user interface for the charge mitigation using chafer is shown Figure S1B. To run this tool, a napari layer associated with the image data is needed, where the charge centres are annotated. This can be set with the button provided. *nlinesaverage, fit length min px* and *fit length max px* are adjustable parameters. *nlinesaverage* sets the number of scanning lines to use to calculate background signal before optimising sigmoid curves. *fit length min px* sets the minimum data points (pixels) next to the annotated artifact that are needed to attempt filtering. Similarly, *fit length max px*, sets the maximum number of points to be used, so that if data along the line extends further, it will be ignored. If this value is too large, more data will be used during the optimization but it may slow calculation significantly.

The resolution estimation tab is shown in Figure S1C. This tool works on two-dimensional data only if data is volumetric, it will calculate in the visualiser selected layer. The calculation splits the image in square tiles with size in pixels being the adjustable parameter *tile size*. The parameter *pixel size* (*nm*), is needed to get accurate results in the statistical information resulting from the FRC calculation.

**Figure S2.**
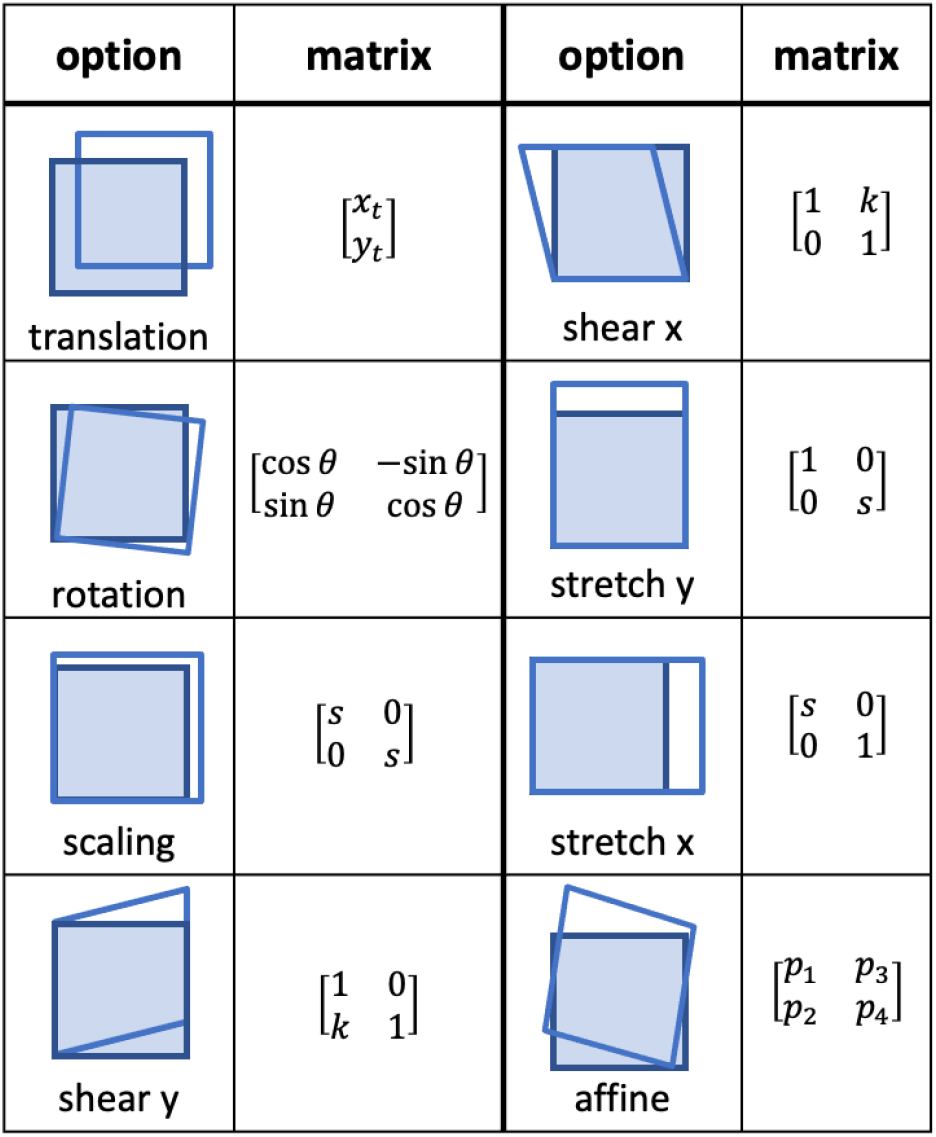
Transformation options available in Okapi-EM as well as their mathematical representations.

**Figure S3.**
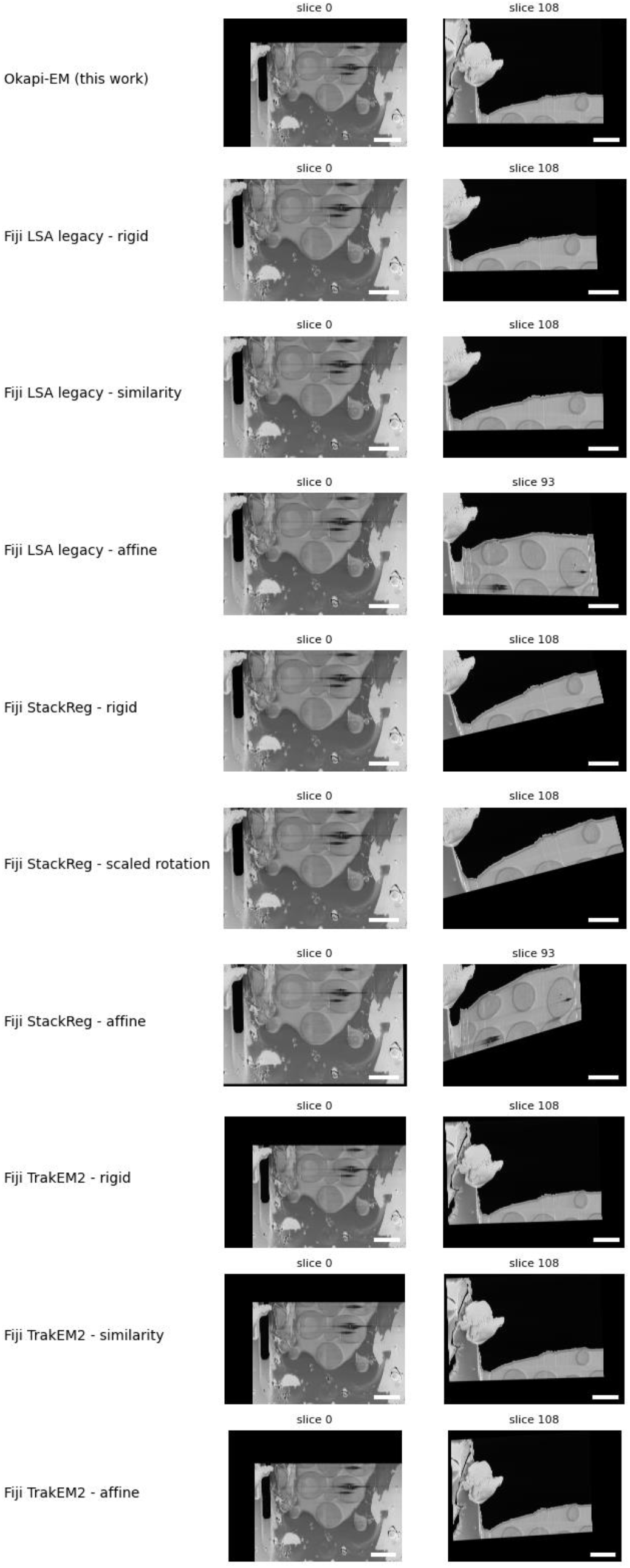
Comparison between different stack alignment algorithms. Various stack alignment algorithms were applied to the same yeast cryogenic serial pFIB/SEM dataset (EMPIAR XXXX). The first and last slice of the stack are presented side-by-side as a means to highlight accumulated matrix transforms that result in excessive rotation in images, which may not reflect the reality of the experiment. All scale bars correspond to 3 μm.

Some important observations to note from the stack alignment results above:

- In most non-okapi alignment algorithms the last image slice is rotated compared to the first slice. StackReg alignments generally result in excessive image rotation.
- Some LSA-Fiji alignments crop slices toward the end (rather than expanding XY image dimensions), which can result in data loss.
- Algorithms *Fiji Legacy - affine* and *Fiji StackReg - affine* did not complete alignment of the whole stack due to errors.

**Figure S4.**
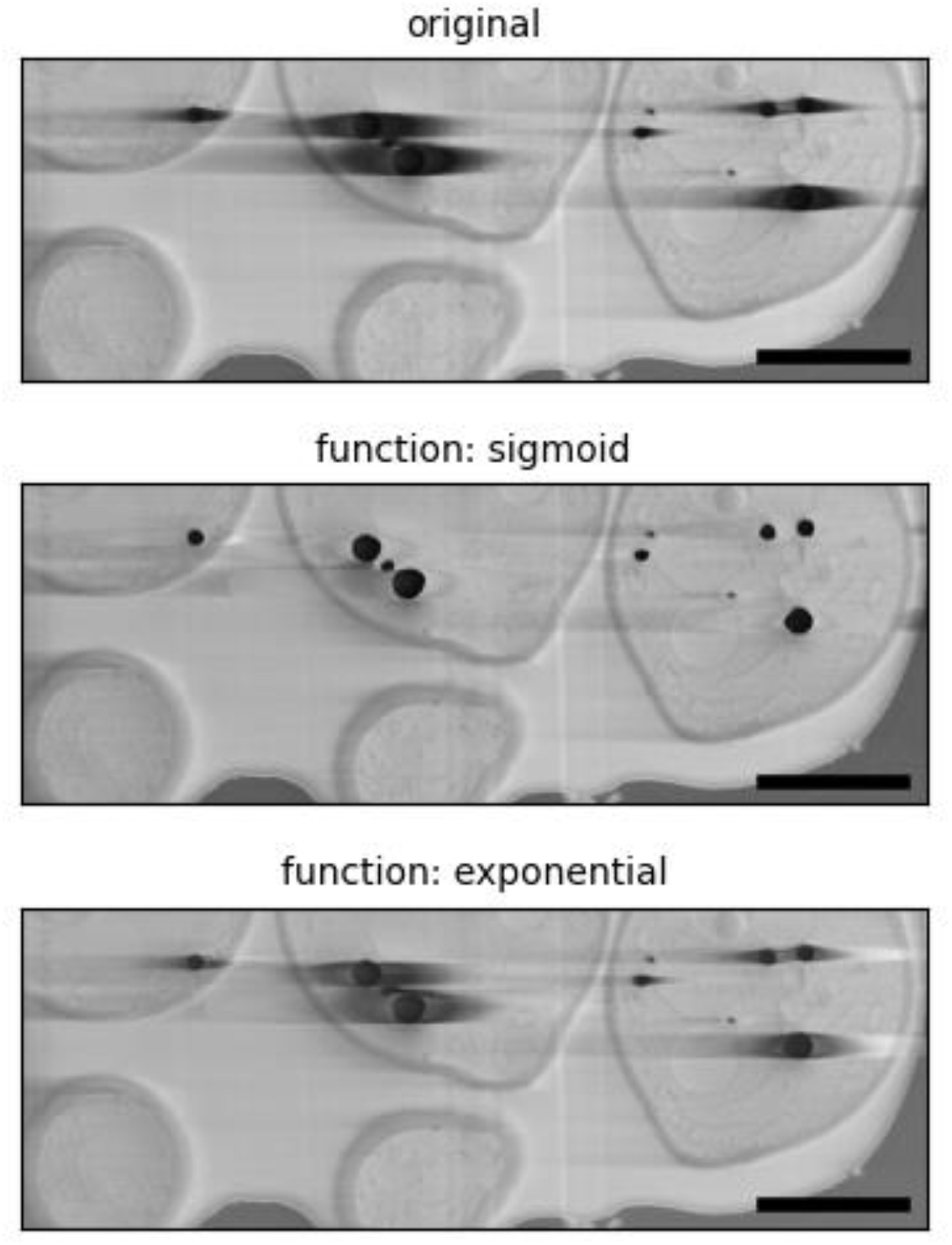
Comparison of results obtained when sigmoid and exponential functions were used for mitigating the charge artifacts using a yeast sample (EMPIAR XXXX). In all calculations, the parameters nlinesaverage, fit length min px and fit length max px used were the default 20, 50 and 700, respectively. Scale bars represent 2 μm.

Visually, the sigmoid filtering gives better results when compared with the exponential method. The exponential filter eliminates some of the tails but the suppression of the artifacts is not as extensive as the sigmoid, and some charging artifacts exhibit additional brightening of the tails on the right side. Chafer tool in Okapi-EM uses the sigmoid implementation of the filter. A gaussian function was also tested and it only successful optimizes with *scipy.optimize* with excessive manual adjustments on a per-row basis. Generally, less than 1% of the optimizations succeeds in a full image, making it inconvenient to use or implement.

As mentioned in the main text, the standard deviation of signal intensity with the charging artifacts masked can be used as a measure of quality of this filter performance, with lower values indicating better removal of the charge artefact tails. For each of the images in Figure S4, from top to bottom, the values obtained are as follow: 23.8, 18.2, 22.4, with the sigmoid filtering presenting a lower value consistent with the visual observations described here.

**Figure S5.**
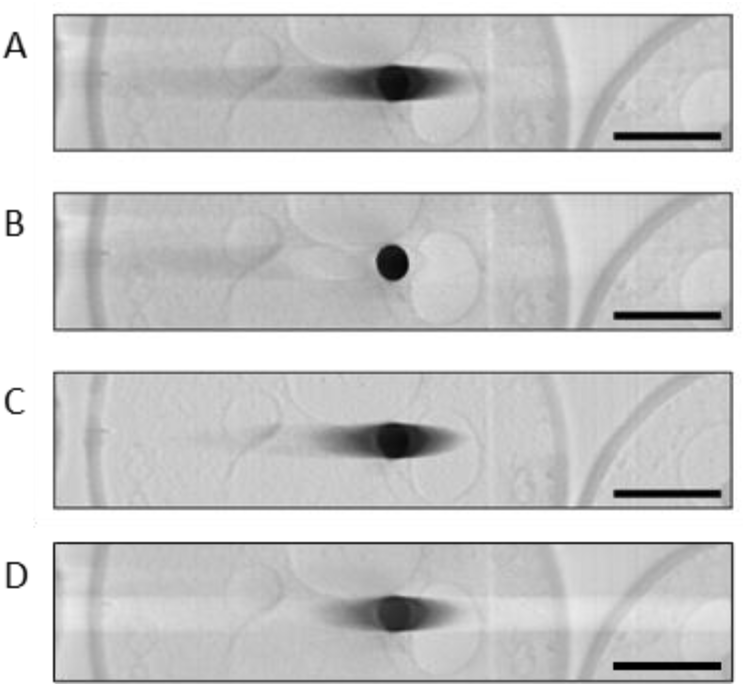
Comparison between charge artefact mitigation filters (A) SEM image of a lipid charge artifact in a yeast sample (EMPIAR XXXX); (B) filtered using chafer in Okapi-EM, (D) using the method described in Spehner et al. (4), and (C) filtered using an FFT bandpass filter in Fiji-ImageJ. All scale bars represent 1 μm.

The charge artifact suppression algorithm presented in Ref (4) by Spehner et al. and further detailed in the “Appendix A. Supplementary data”, was tested with the charging artifacts in our images, as described in the publication. However only minor suppression is observed (Figure S5C), mainly further away from the charging centre.

The FFT bandpass filter in Fiji ImageJ was also tested and it did not result in a substantial charge suppression (Figure S5D). The filter settings used were:

- structures by size: zero
- stripe suppression: horizontal and tolerance of direction 5%
- no autoscaling of intensities.

**Figure S6.**
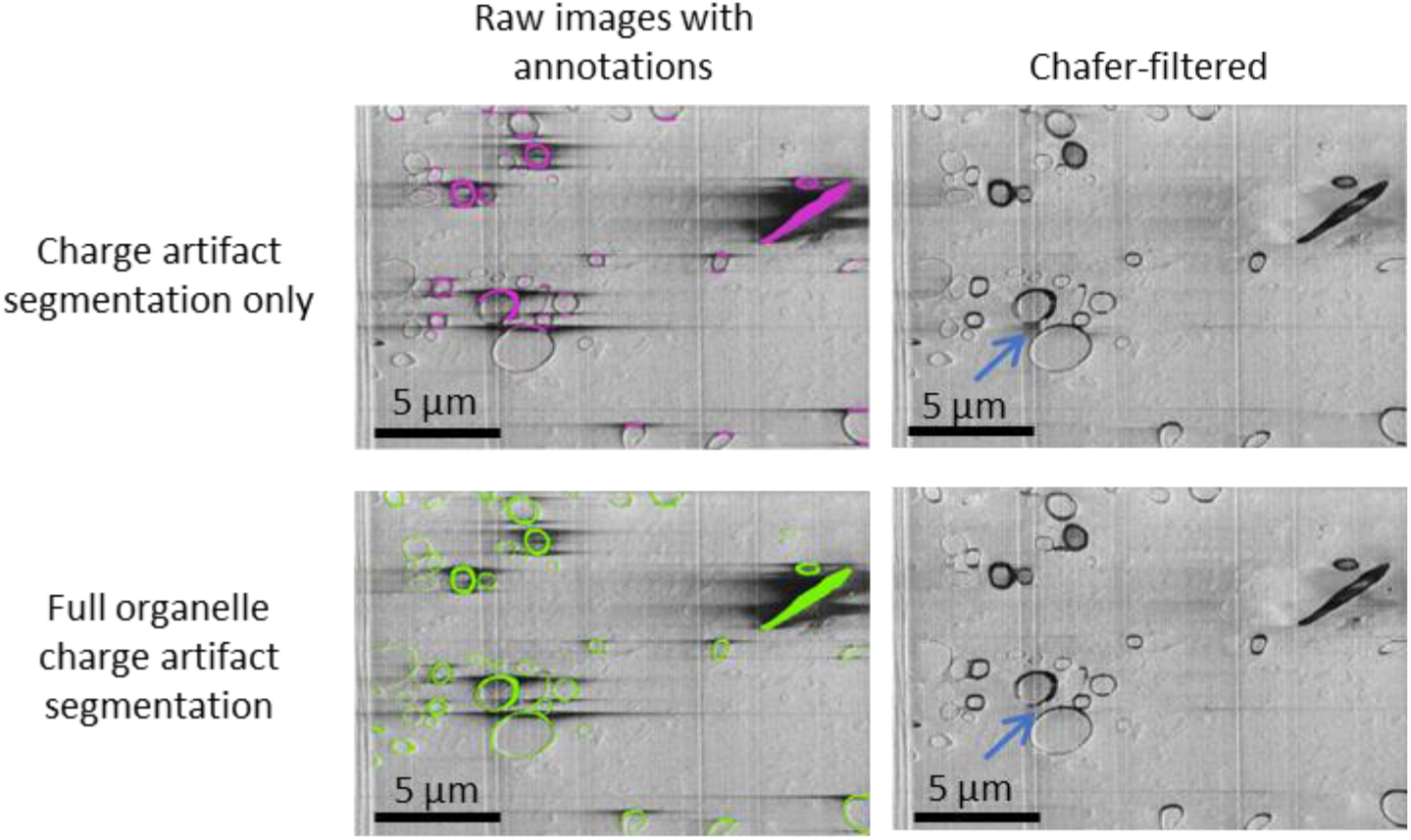
Chafer performance when only charging features are included vs when whole organelles (regardless of charging status) are included. Left, SEM images of mouse brain (EMPIAR XXXX). Left, top, where only charging artifacts have been segmented as indicated in pink; in left, bottom the whole organelle responsible for the emergence of charging artifacts (myeline sheaths) is fully segmented in green. Right column is the result after running the chafer filter using the respective labels on the left. The resulting images are visually similar, with the most significant difference indicated with a blue arrow.

